# In utero exposure to morphine leads to sex-specific behavioral alterations that persist into adulthood in cross-fostered mice

**DOI:** 10.1101/2022.02.28.482336

**Authors:** Vanessa C. Fleites, Patrick S. Markwalter, Keenan Johnson, Mariella De Biasi

**Author notes:** Corresponding Author: Mariella De Biasi.

## Abstract

**Introduction:** The opioid epidemic has seen an increase in drug use among women of reproductive age. It is well established that Opioid Use Disorder (OUD) can have many negative consequences for the health of mothers and their babies, both during pregnancy and after delivery, but our understanding of the impact of fetal opioid exposure on behavior during adolescence and adulthood is less understood. Preclinical studies have unveiled some of the long-term effects of *in utero* morphine exposure primarily using injections as the route of drug delivery. Our study utilized a model for oral, voluntary morphine self-administration to investigate neonate, adolescent, and adult offspring’s behavioral phenotypes and subsequent ethanol misuse liability.

**Methods:** We first validated a paradigm for maternal oral intake of morphine, where female mice became morphine dependent pre-pregnancy, and continued to voluntarily consume morphine in the continuous two-bottle choice (C2BC) paradigm during pregnancy and up to offspring postnatal day 7 (PND 7). Offspring were cross-fostered to a drug-naïve dam at PND 7, to model first and second trimester *in utero* exposure in humans and to mimic the stress associated with NOWS. Bodyweight and ultrasonic vocalizations were assessed to determine alterations in the neonates. Offspring from control and morphine-exposed dams were then tested during adolescence and adulthood in a battery of behavioral tests to assess baseline behavioral phenotypes. We also computed a global behavioral score (GBS) to integrate offspring’s multiple behavioral outcomes into a composite score that could be used to identify potential vulnerable and resilient populations in offspring exposed prenatally to morphine. Offspring that were tested during adolescence were also evaluated during adulthood in the ethanol intermittent 2BC to assess ethanol misuse risk.

**Results:** Using an oral maternal morphine C2BC protocol, we demonstrated that morphine dams display signs of dependence, measured by somatic signs during withdrawal, and voluntarily drink morphine throughout gestation. Neonate cross-fostered offspring display changes in spontaneous activity, body weight, and ultrasonic vocalization parameters. During adolescence, offspring display both increased baseline anxiety-like/compulsive-like behavior, while in adulthood they display increased anxiety-like behavior. No changes were found for baseline physical signs, locomotion, and depressive-like behavior during adolescence or adulthood. In addition, a greater percentage of adult male offspring exposed to maternal morphine fell into moderate and high GBS classifications, signaling a more severe behavioral phenotype, compared to male control offspring. These effects were not observed in adult female offspring exposed to morphine *in utero*. Additionally, male adult offspring exposed to maternal morphine reduced their 2-hour ethanol intake in the intermittent two-bottle choice (I2BC) paradigm, although no changes in 24-hour ethanol intake and preference were found. No changes were observed in female offspring of morphine-exposed dams.

**Conclusion:** Overall, maternal morphine exposure leads to sex-specific changes in neonate, adolescent, and adult behavior, including ethanol intake.

## 1 Introduction

Opioid Use Disorder (OUD) is a major public health concern. The 2019 National Survey on Drug Use and Health reported that 3.7% of individuals aged 12 and older, including women of childbearing age, have misused prescription and/or illicit opioids (Lipari & Park-Lee, 2020). In addition, a 2015-2016 study showed that one third of pregnant women used opioids (St. Marie, Coleman, Vignato, Arndt, & Segre, 2020). Continued opioid use in pregnant women can lead to serious maternal, fetal, and neonate complications and, in extreme cases, can lead to death (Leyenaar et al., 2021; Ostrea, Ostrea, & Simpson, 1997).

Some newborns prenatally exposed to opioids display a series of symptoms categorized as Neonatal Opioid Withdrawal Syndrome (NOWS). A study of maternal-infant dyads prenatally exposed to opioids reported that roughly 30-60% of newborns were diagnosed with NOWS (Leyenaar et al., 2021; Skumlien, Ibsen, Kesmodel, & Nygaard, 2020), which suggests that a proportion of newborns exposed to opioids *in utero* may develop a phenotype severe enough to require pharmacological and/or non-pharmacological interventions. Characteristic manifestations of NOWS include decreased body weight, high-pitched crying, irritability, tremors, and an inability to be soothed, among many others (Piccotti et al., 2019; Weller, Crist, Reiner, Doyle, & Berrettini, 2020). Reviews and meta-analyses of clinical data have reported infant-adolescent outcomes associated with *in utero* opioid exposure, including lower scores in neurocognitive and developmental assessments, decreased motor skills, and increased hyperactivity and aggression (Maguire et al., 2016; Minnes, Lang, & Singer, 2011; Nygaard, Slinning, Moe, & Walhovd, 2017; Weller et al., 2020; Yeoh et al., 2019). However, other studies report variability in outcomes among children exposed prenatally to opioids, leading to potential differences in vulnerability and resiliency in these individuals (Labella, Eiden, Tabachnick, Sellers, & Dozier, 2021; Sarfi, Eikemo, Welle-Strand, Muller, & Lehmann, 2021). In addition, the long-term effects of maternal opioid use on human offspring are not fully understood, including the vulnerability to develop a psychiatric disease and the risk of drug misuse.

Morphine is a mu-opioid receptor (MOR) agonist and an active metabolite of heroin, so it is commonly used in studies investigating the effects of acute and chronic opioid exposure. Morphine is also the standard of care used for acute pain, and is used to sedate patients pre- and post-operatively, including mothers and newborns (Doleman et al., 2018; Lugo & Kern, 2002). Preclinical models have been used extensively to study the long-term effects of maternal morphine exposure. Several rodent studies have reported the effects of prenatal and perinatal morphine exposure on offspring outcomes, including body weight, mortality, and organ size (Ahmadalipour, Ghodrati-Jaldbakhan, Samaei, & Rashidy-Pour, 2018; Chiang, Hung, & Ho, 2014; Eriksson & Rönnbäck, 1989; Glick, Strumpf, & Zimmerberg, 1977; Klausz et al., 2011; Ramsey, Niesink, & Van Ree, 1993; Shen et al., 2016; Siddiqui, Haq, & Shah, 1997; Sobor et al., 2010; Tan et al., 2015; Timár et al., 2010). However, very few of them have examined other aspects of NOWS, such as high-pitched crying. Similar to the studies in humans, preclinical models of *in utero* morphine exposure report discrepant offspring outcomes. Many factors could contribute to such variability, including differences in maternal opioid exposure paradigms, including length of exposure and dose used, and whether dams experienced varying levels of gestational stress, like being shipped while pregnant or receiving daily injections. To date, no published study has used an oral two-bottle choice (2BC) morphine self-administration protocol for the maternal dam exposure, which minimizes any confounds associated to stress.

The preclinical literature has also shown conflicting results on whether parental morphine causes alterations in behavior related to psychiatric disorders, including anxiety-like, compulsive-like, and depressive-like behavior. For example, in one study adult rodents exposed to morphine *in utero* displayed decreased anxiety in both the Elevated Plus Maze (EPM) and the Light-Dark Box (Tan et al., 2015). In contrast, a study where morphine was given between gestation day (GD) 1 and postnatal day (PND) 21 found no differences in offspring’s anxiety-like behavior in the EPM (Klausz et al., 2011). Additionally, most studies have only examined a few offspring behaviors at a time, and to date, no study has evaluated behaviors with a comprehensive approach, to better define baseline phenotypes of adolescent and adult offspring after maternal opioid exposure.

Parental substance use disorder (SUD) and prenatal drug exposure is associated with a myriad of negative outcomes in offspring, including vulnerability to develop a SUD (Dodge, Jacobson, & Jacobson, 2019). Negative outcomes in the offspring depend on the timing of the parental exposure to drugs and/or other external factors (Betcher et al., 2019; Biederman, Faraone, Monuteaux, & Feighner, 2000; Dodge et al., 2019; Glantz & Chambers, 2006; Madras et al., 2019; Peleg-Oren & Teichman, 2006; Tarter et al., 2020). Pre- and perinatal opioid exposure can alter offspring’s predisposition to future drug use, including risk for the same drug-class from maternal exposure (i.e. morphine) or cross-tolerance to other drugs (i.e. cocaine and methamphetamine) (Chiang et al., 2014; Gagin, Kook, Cohen, & Shavit, 1997; Glick et al., 1977; Ramsey et al., 1993; Shen et al., 2016; Timár et al., 2010; Vousooghi et al., 2018; L. Y. Wu, Chen, Tao, & Huang, 2009). To our knowledge, no preclinical study has examined potential changes in alcohol intake and preference in offspring exposed to *in utero* morphine, even though alcohol is the most commonly used drug according to the 2019 National Survey on Drug Use and Health (Lipari & Park-Lee, 2020). One clinical longitudinal study reported that a significantly higher proportion of adults whose mothers used heroin during pregnancy misused alcohol during their lifetime (Nygaard, Slinning, Moe, Fjell, & Walhovd, 2020). Furthermore, there is well-documented evidence for opioid-ethanol interactions, warranting further investigation into how pre- and perinatal opioid exposure affects alcohol use risk (Arias & Kranzler, 2008; Gianoulakis, 2001; Gianoulakis et al., 1989; Job et al., 2007).

In the present study, we utilized a translational maternal morphine paradigm, where dependent dams orally self-administered morphine in a 2BC paradigm throughout pregnancy and during the first postnatal week to model morphine exposure during early- and mid-gestation in humans (Richard & Flamant, 2018; Semple, Blomgren, Gimlin, Ferriero, & Noble-Haeusslein, 2013). We examined offspring in a battery of behaviors throughout adolescence and adulthood to compile a comprehensive behavioral score. After adolescent testing, offspring were monitored for alcohol oral self-administration during adulthood, revealing sex-specific changes in ethanol intake.

## 2 Materials and Methods

### 2.1 Research subjects

C57BL/6J mice of both sexes (IMSR Cat# JAX:000664, RRID:IMSR_JAX:000664) were given access to housing enrichment and *ad libitum* food and water (Labdiet, 5053, PMI, Brentwood, MO). The animals were maintained on a reverse 12-hr light/12-hr dark cycle (lights “off” at 10:00 AM) and housed in a temperature- and humidity-controlled room (65-75 °F, 40-60% relative humidity). All experiments were conducted during animals’ active phase (10:00 AM – 8:00 PM). For all drinking experiments, an empty control cage was set up with two bottles that were weighed daily to account for fluid leakage due to cage and bottle handling. For all behavioral tests, animals were habituated to the testing room and light conditions at least 30 minutes prior to the start of the test. A digital light meter was used to measure luminosity in the room for each test, reported as lux. All experiments were conducted with the approval of the Institutional Animal Care and Use Committee (IACUC) at the University of Pennsylvania.

### 2.2 Maternal Drug Exposure

Female mice were habituated to being single-housed and to being exposed to one bottle of 0.2% saccharin (Sigma, St. Louis, MO) in filtered water for one week (Figure 1a-Habituation). After the one-week habituation, female mice were observed for 30 minutes in their home cage for baseline physical signs (room luminosity at 2 lux). Signs commonly reported for morphine withdrawal were included in the analysis: jumping, wet dog shakes, head nods/shakes, teeth chattering, diarrhea, and writhing (Muldoon et al., 2014; Pinelli & Trivulzio, 1997). Female mice were then separated into an experimental (morphine + saccharin) group, referred to as “morphine dam”, and a control (saccharin) group, referred to as “control dam”. Groups were created considering the baseline physical signs, to ensure that both groups had on average a similar total number of physical signs to start. The morphine dams were given one bottle of 0.1 mg/mL free-base morphine (Morphine Sulfate, Spectrum Chemical MFG. Corp, New Brunswick, NJ) in 0.2% saccharin for four days and then the solution was escalated to 0.2 mg/mL free-base morphine + saccharin for three days (Figure 1a). Bottles were weighed daily, and their position was alternated to avoid side preference. Morphine intake is reported as weekly averages in mg/kg/day. To confirm dependence, female mice were tested for spontaneous signs of withdrawal 8 hours after the morphine bottle was replaced with a saccharin-only bottle, at the end of the one-week forced morphine exposure. Control female mice were maintained on one bottle of 0.2% saccharin throughout this time and observed for physical signs.

**Figure 1:**
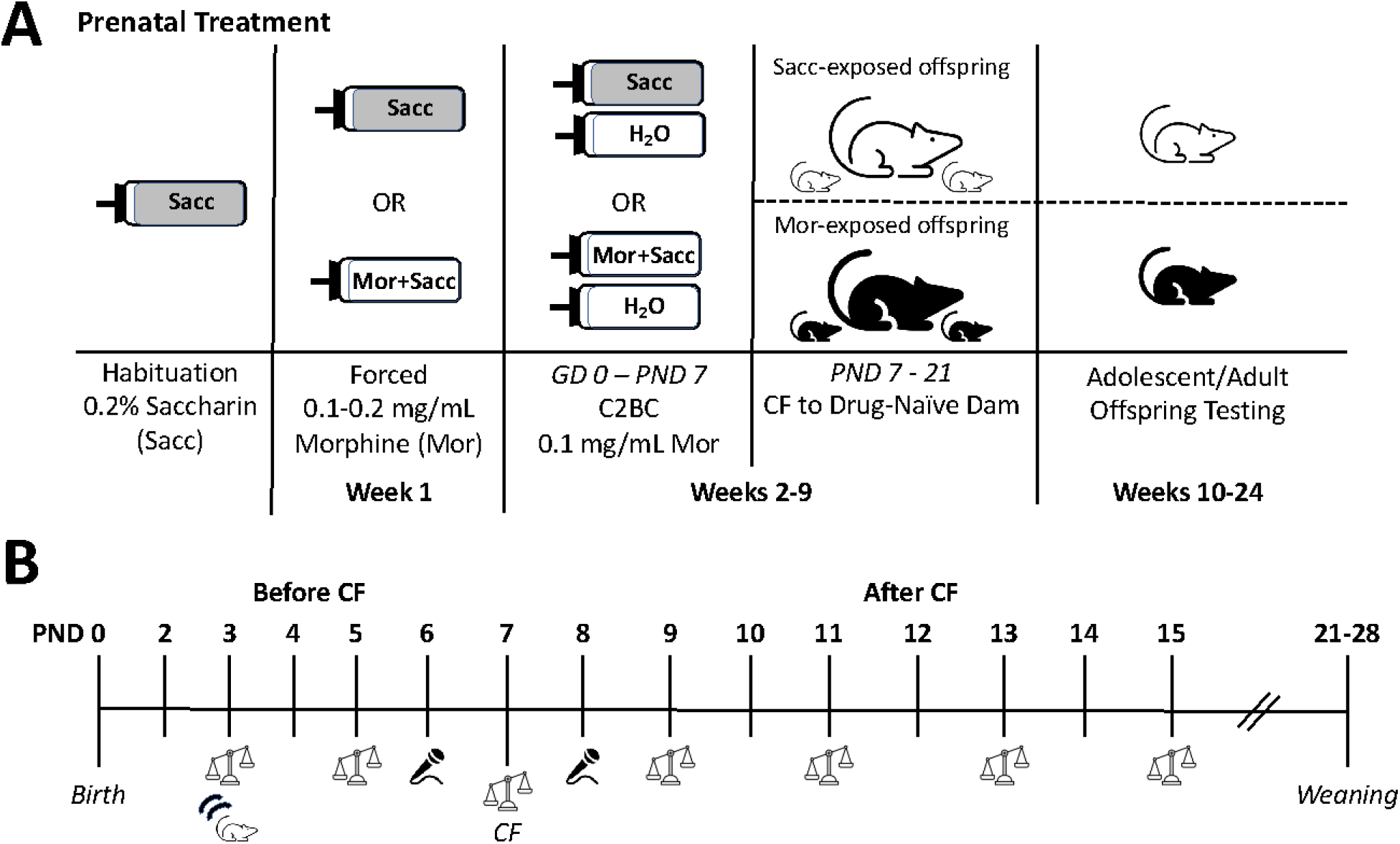
Experimental schemes for maternal morphine exposure and offspring behavioral evaluation. **A**. After a week of habituation to 0.2% saccharin, female mice drank from a single bottle containing either 0.2% saccharin or 0.2 % saccharin + morphine. Mice were then transitioned to a C2BC paradigm that lasted throughout mating, gestation, and the first week after delivery. On PND 7, offspring were cross-fostered (CF) to drug-naïve dams and were subsequently tested in various behavioral paradigms during both adolescence and adulthood. **B**. Evaluation of offspring behavior prior to weaning (PND 0 – 28) was conducted before and after cross-fostering and included observation of spontaneous activity (moving mouse icon) on PND 3, recording of ultrasonic vocalizations (microphone icon) on PND 6 and PND 8, and measurement of body weight (scale icon) on PND 3, 5, 7, 9, 11, 13, 15. GD = Gestation Day; Mor= morphine; Sacc=saccharin; H2O= water; PND= Postnatal Day; C2BC= continuous two-bottle choice

The female mice were then transitioned to the continuous two-bottle choice (C2BC) phase the day after somatic signs testing (Figure 1a). Mice in the morphine dam group received one bottle containing 0.1 mg/mL free-base morphine in 0.2% saccharin, and a bottle containing filtered water. Mice in the control dam group received one bottle of 0.2% saccharin solution and a bottle of filtered water. After one week of the C2BC, mice from the morphine dam group were evaluated for spontaneous physical signs of withdrawal 8 hours after removal of the morphine bottle, as described above. During week two of the C2BC paradigm, mice were evaluated again for physical signs while drug sated. Mice from the control dam group were evaluated for physical signs on the same day as the mice from the morphine dam group.

Because we were interested in investigating only the effects of maternal opioid use, female mice were placed daily in the cage of single-housed, drug-naïve males to mate for approximately five hours/day, and were then returned to their home cage to continue their C2BC paradigm throughout gestation. Pregnancy was confirmed by a substantial increase in body weight after one week. The day at which pups were found is referred to as postnatal day 0 (PND 0). Offspring were allowed to lactate from their respective dam until PND 7 (Figure 1b). On PND 7, offspring were cross-fostered to an experienced drug-naïve dam, who had her own litter removed at the time of cross-fostering. We chose to cross-foster offspring on PND 7, because the first postnatal week is roughly equivalent to the human second trimester/early third trimester, with regards to brain maturation (Barr, McPhie-Lalmansingh, Perez, & Riley, 2011; Ross, Graham, Money, & Stanwood, 2015; Semple et al., 2013). In addition, cross-fostering would allow us to evaluate early neonate outcomes after acute morphine withdrawal.

On PND 3, modified spontaneous physical signs from Barr *et al*. (2011) were recorded for 2 pups per litter (6-7 litters/ dam treatment), which included audible cries, rolling over, and full 360° body rotation. Each pup was removed from its cage and placed on a paper towel, under red light to minimize stress. Pups were observed for five minutes for spontaneous signs, and then immediately returned to their respective litter. Offspring were weighed on PND 3 and weighed every other day (odd days) thereafter until PND 15 (Figure 1b). Litter averages are shown for body weight data, since individual pups were not tattooed to keep track of ID. All male and female offspring cross-fostered from both control dams and morphine dams were weaned between PND 21-28 and used for neonate, adolescent, and adult testing.

### 2.3 Ultrasonic Vocalizations

Evaluation of neonate offspring ultrasonic vocalizations (USVs) from both treatment groups began on PND 2 and ended on PND 12 (Figure 1b). USVs were recorded on PND 2, 4, and 6 to evaluate changes related to lactation from control dams and morphine dams. USVs were also recorded on PND 8, 10, and 12 to evaluate changes associated with cross-fostering on PND 7 and potentially, morphine withdrawal, as observed in human newborns that experience NOWS. We were specifically interested in USV parameters before and after cross-fostering, so only data for PND 6 and PND 8 are reported (Figure 1b).

Individual pups were transferred into a small container with bedding and placed in an enclosed Styrofoam box with an ultrasonic microphone (Dodotronic Ultramic, Dodotronic, Castel Gandolfo, Italy) inserted on top. The microphone was connected to a laptop that was running Raven Pro 1.6.0 (Raven, RRID:SCR_016190, Center for Conservation Bioacoustics, Cornell Lab of Ornithology, Ithaca, NY) to save and analyze the file as a 120-megabyte.wav file. Offspring USVs were recorded for five minutes. An average of three pups per litter was recorded on the assigned even-day PND.

USVs from each sound file were visualized using Raven software’s spectrogram and were manually selected by an experimenter blinded to the treatment groups to avoid bias. The selections made were automatically incorporated into an aggregate Selection Table produced by Raven that gave various measurements for each call selected, including number of syllables, low frequency (Hz), high frequency (Hz), delta time (s), delta frequency (Hz), and center frequency (Hz). In Raven, delta time is described as the average difference between the start and end time of each call in the sound file. Delta frequency is defined as the average difference between the maximum and minimum frequency of each call in the sound file. Center frequency is defined as the average middle frequency of each call in the sound file.

We calculated the average of each measurement in the Selection Table for each pup’s PND USV 5-minute file, to provide an average for a specific USV parameter for each individual pup within a litter. To compare litters across PNDs (7-9 litters/ dam treatment), the values for USV measurements for pups within a litter were averaged for a given PND based on their treatment groups, giving us a “litter average”.

### 2.4 Adolescent baseline behavioral tests

Offspring were habituated to handling at least five days before adolescent testing started. Testing during adolescence occurred between PND 28 and PND 49.

To assess baseline physical signs, adolescent offspring were observed for shaking, scratching, grooming, and teeth chattering as described before (E. Perez, Quijano-Cardé, & De Biasi, 2015; Ramiro Salas, Main, Gangitano, & De Biasi, 2007). Offspring were placed in a novel cage with clean corncob bedding (room lux: 2) and observed for 20 minutes (6-9 litters examined/dam treatment). The same cages were then used for the marble burying test (MBT) to assess anxiety-like/compulsive-like behavior as previously described (Njung’e & Handley, 1991; E. E. Perez & De Biasi, 2015). Briefly, cages were filled with 5 centimeters of bedding and 20 marbles evenly spaced on top. Offspring were left undisturbed for 30 minutes (room lux: 2) and the number of marbles buried (fully buried or at least 2/3 buried) was recorded.

At least 48 hours after the MBT, offspring were tested in the Open Field Arena (OFA) test. The OFA consisted of a white plexiglass squared platform (40 centimeters by 40 centimeters) with walls (Gangitano, Salas, Teng, Perez, & De Biasi, 2009; Ramiro Salas, Pieri, Fung, Dani, & De Biasi, 2003). The OFA was divided into a center zone (20 cm by 20 cm) and a surround zone (10 cm from wall all around OFA). The average center zone luminosity was 4 lux, while the corner surround zone luminosity was 2 lux. Offspring were placed at the center of the OFA and allowed to freely explore for 30 minutes while being recorded with ANYMAZE software (Stoeling Co, Wood Dale, IL). Locomotion and anxiety-like behavior were assessed by measuring the average total distance travelled (m) and center distance ratio (distance travelled in center zone (m)/total distance travelled (m)), respectively.

At least 48 hours after the OFA, offspring were tested in the Elevated Plus Maze (EPM), as previously described (E. E. Perez & De Biasi, 2015; Ramiro Salas et al., 2003). The luminosity used for the open arms was 4 lux, and that for the closed arms was about 1 lux. Animals were placed in the center zone of the EPM and allowed to freely explore for 10 minutes. Average time spent in the open arms (s) and open arm entry ratio (open arm entries/ open arm entries + closed arm entries) were reported to evaluate anxiety-like behavior.

At least 48 hours after the EPM test, offspring were tested in the Tail Suspension Test (TST) to measure depressive-like behavior as previously described (Gangitano et al., 2009; R. Salas et al., 2008). The luminosity of the area under the tail suspension apparatus was about 4 lux. Tape was used to hold the tail onto the TST apparatus, and the animal was hung upside down for six minutes. Average time spent immobile (s) was reported.

#### Global Behavioral Score Classification

Six behavioral measures were used to calculate global baseline behavioral scores (GBS) in offspring. The measures include: (1) physical signs, calculated as the total number of physical signs, (2) anxiety-like/compulsive-like behavior, calculated as total number of marbles buried in MBT, (3) locomotion, calculated as total distance travelled in OFA, (4) anxiety-like behavior in OFA, calculated as center distance ratio, (5) anxiety-like behavior in EPM, calculated as open arm entry ratio, (6) depressive-like behavior, calculated as total immobility time in the TST. The measures used for the GBS were chosen *a priori* based on our hypothesis that offspring from morphine-exposed dams would display increased baseline physical signs, compulsive-like behavior, anxiety-like behavior, depressive-like behavior, and hyperlocomotion. Because our hypothesis included changes in locomotion, we used measures of anxiety-like behavior that incorporated locomotion in the measure, instead of using time in a zone (i.e. if locomotion is changed then that might influence time spent in a particular zone and affect anxiety-like measures).

Similar to O’Neal *et al*. (2020) and Quijano Cardé *et al*. (2022), z-scores were calculated for each behavioral measure. Briefly, the group mean (*µ*^1^) for each behavioral measure was subtracted from the raw individual value (*x*^1^) for each offspring for that behavior, and then divided by the group standard deviation, 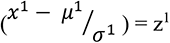. The z-score was then multiplied by the direction (+1 or -1) for that behavioral measure, to indicate worst behavioral outcome. For example, for center distance ratio in the OFA and open arm entry ratio in the EPM, the lower the raw value, the more anxiety-like behavior the offspring displays, so the z-score for both of these behavioral measures would be multiplied by -1 to correct for the direction. Conversely, higher raw values for marbles buried indicate increased compulsive-like behavior, so the z-score is multiplied by +1 to reflect a worst behavioral outcome. Individual z-scores for each offspring were added to obtain a global behavioral score for that subject 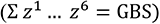. Only offspring with raw data for all behavioral measures were used for this analysis.

### 2.5 Adult ethanol intermittent two-bottle choice (I2BC) paradigm

Mice that were previously analyzed for adolescent baseline behaviors were examined for ethanol drinking behavior during adulthood using the ethanol I2BC, as previously described (Carnicella, Ron, & Barak, 2014; Hwa et al., 2011; Quijano Cardé & De Biasi, 2022; Quijano Cardé, Perez, Feinn, Kranzler, & De Biasi, 2021). Offspring (at least 2 months of age) were habituated to being single-housed and were exposed to two 50 mL bottles of filtered water for at least one week in the home cage. Afterwards, mice were given 24-hour access to a bottle of ethanol and a bottle of water on Mondays, Wednesdays, and Fridays. On alternating days, mice were presented with two bottles containing filtered water. During week 1 or the ‘Acquisition’ phase, mice were habituated to ethanol by receiving increasing concentrations of ethanol: 3% (Monday), 6% (Wednesday), and 10% (v/v) ethanol (Friday). During weeks 2-5 of the ‘Maintenance’ phase, mice were given one bottle of 20% ethanol (v/v) and one bottle of water. Mice were then transitioned to a ‘Fading’ phase of the experiment to determine if mice would drink more to maintain the same ethanol dose they received on week 6 (20% ethanol) even when the ethanol concentration was progressively decreased during the subsequent weeks. During weeks 7-10, mice were given decreasing concentrations of ethanol each week (15%, 10%, 6%, and 3% ethanol). Presentation of the ethanol bottle occurred three hours after ‘lights off’ (1:00 PM), and 2- and 24-hour ethanol consumption (g/kg/day) and preference [(ethanol ml intake/total ml fluid intake)*100%] were measured. All mice were weighed weekly. All ethanol solutions were made in filtered water (v/v) using 190-proof ethanol (Decon Laboratories Inc., King of Prussia, PA).

### 2.6 Adult baseline behavioral tests

In a separate cohort, adult (at least two months of age) group-housed offspring from control and morphine-treated dams were habituated to handling. Physical signs, MBT, OFA, EPM, and TST were examined at least 24-hours apart. To assess baseline physical signs, adult offspring were observed for jumping, shaking, scratching, grooming, and teeth chattering. Offspring were also assessed for baseline behaviors in the MBT, OFA, EPM, and TST, like described in the previous sections.

### 2.7 Statistical analyses

Data were analyzed using Graphpad PRISM 9 and are expressed as mean +/-standard error of the mean (SEM). Litter averages are shown for neonate body weight and USV data, while individual data points are shown for dam, adolescent, and adult data. Dam and neonate outcomes were analyzed using paired t-test, t-test, or repeated measures (RM) one-way ANOVA, when appropriate. Tukey post-hoc analysis was used as recommended. Adolescent offspring data were first analyzed using a two-way ANOVA to investigate the potential effect of sex and/or dam treatment, and the interaction between the two variables. Since no effect of sex was observed, a one-way ANOVA was used for analysis of adolescent and adult offspring behavioral data. GBS classifications for adolescent and adult offspring data were analyzed using an outcome versus expected chi-square test, where the control offspring percentages for each classification (‘low’, ‘moderate’, ‘high’) were used as the ‘expected’ percentages to compare to percentages obtained from offspring from morphine-exposed dams. A RM three-way ANOVA for ethanol I2BC drinking data revealed a main effect of sex, so males and females were analyzed separately using a RM two-way ANOVA with Sidak post-hoc analysis. For the I2BC experiment, the ‘Acquisition’, ‘Maintenance’, and ‘Fading’ phases were analyzed separately. For datasets missing values at certain experimental timepoints, a mixed effects model with a Sidak post-hoc test was performed. A *p*-value of <0.05 was considered statistically significant. ROUT (Q = 1%) was used to remove significant outliers.

## 3 Results

### 3.1 Validation of a maternal morphine exposure model in mice

We developed a paradigm to model opioid use in humans and ultimately investigate its effects on offspring, by using an oral morphine self-administration protocol in female pregnant mice (Figure 1a). Since human mothers who are opioid-dependent begin drug use before pregnancy, we first established a paradigm where breeding-age female mice would become dependent on morphine. To create initial opioid dependence, mice were given one bottle of escalating concentrations of morphine (0.1 mg/mL – 0.2 mg/mL), which led to increased morphine intake (paired t-test; *t* = 5.896, *df* = 10, *p* = 0.002; Figure 2a). Under this treatment paradigm, female mice displayed increased spontaneous physical signs of withdrawal 8 hours after the removal of the morphine bottle, compared to their pre-treatment baseline (paired t-test; *t* = 6.835, *df* = 9, *p* = <0.0001; Figure 2b). Mice were then transitioned to a continuous two-bottle choice (C2BC) paradigm, where they received one bottle of morphine in saccharin water and one bottle of water. Based on previous 2BC morphine protocols used in the field, saccharin was added only to the morphine bottle because morphine salt is perceived as bitter and we wanted to limit the variability of the dose of morphine consumed between dams (Belknap, 1990; Belknap, Crabbe, Riggan, & O’Toole, 1993; Ferraro et al., 2005). After one week in the morphine C2BC paradigm (week 2 of the paradigm), mice drank on average 37 mg/kg morphine solution (Figure 2c). Mice also displayed increased spontaneous physical signs of withdrawal 8 hours after the removal of the morphine bottle, compared to when they were morphine-sated in the C2BC (paired t-test; *t* = 4.315, *df* = 9, *p* = 0.0019; Figure 2d), and compared to control female mice that received drug-free sweetened fluid (t-test; *t* = 3.484, *df* = 20, *p* = 0.0023; Figure 2e).

**Figure 2:**
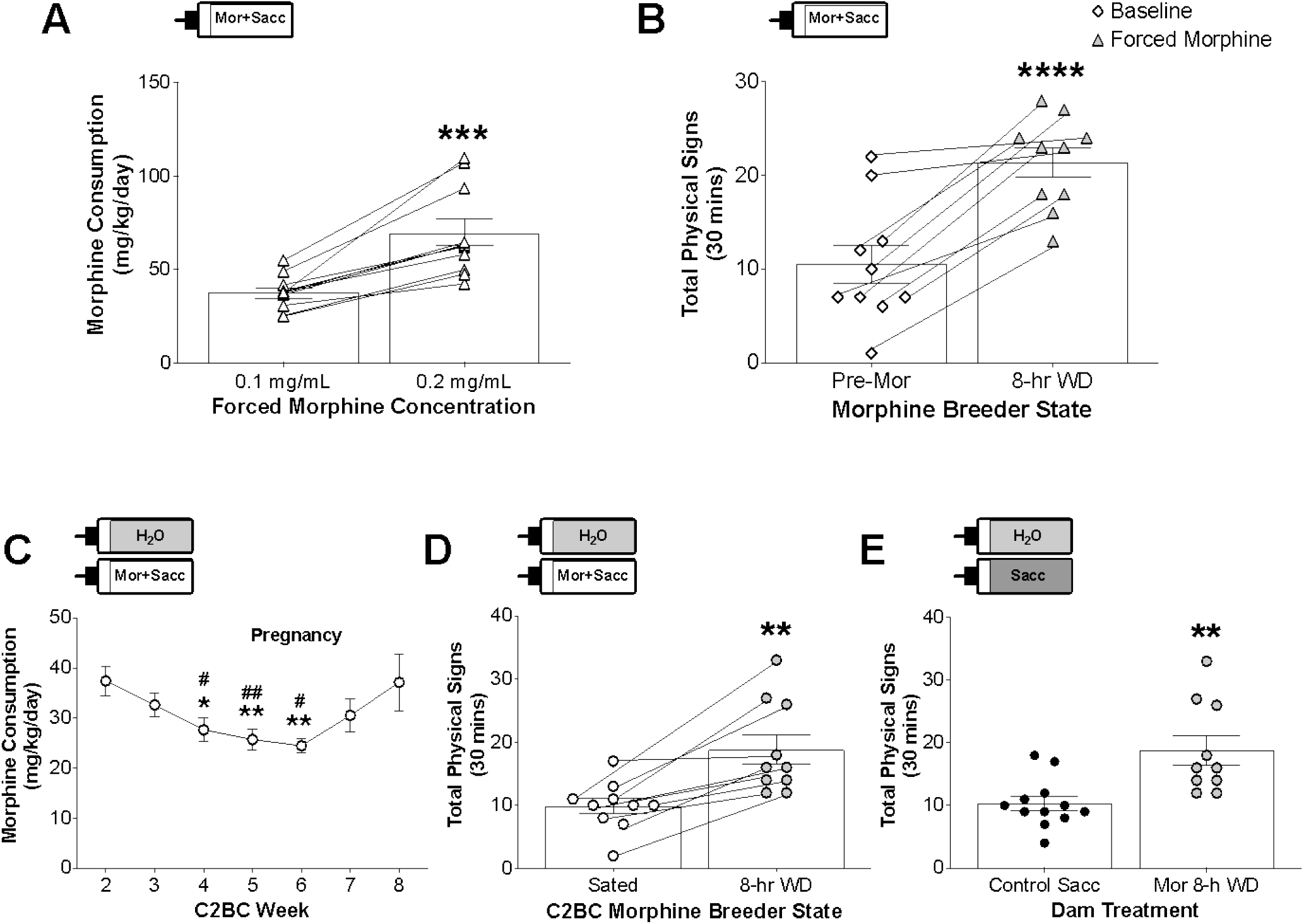
Validation of the paradigm for maternal morphine exposure. (**A & B**) Morphine consumption and physical signs measured in female breeders while having access to a single bottle containing morphine (forced morphine exposure). **A**. Forced morphine intake during the initial phase of treatment, when morphine dams’ solution is ramped up from 0.1 mg/mL morphine to 0.2 mg/mL morphine, respectively (n=11). **B**. Total physical signs in morphine-exposed dams at baseline (pre-treatment), and 8 hours after withdrawal from forced morphine exposure (n=10). (**C-E**) Morphine intake and physical signs during the C2BC paradigm. **C**. Morphine intake in the C2BC paradigm during weeks 2-8. (n=10-11/week) * p<0.05, **p<0.01 compared to Week 2; # p<0.05, ## p<0.01 compared to Week 3. **D**. Total physical signs for morphine-exposed dams while morphine sated in the C2BC paradigm and 8 hours after withdrawal from morphine (n=10). **E**. Comparison of total physical signs in control, saccharin-drinking dams and morphine-drinking dams 8 hours after morphine withdrawal in the C2BC paradigm (n=12,10). ** p<0.01, ***p<0.001, **** p<0.0001; Mor= morphine, WD= withdrawal, C2BC= continuous two-bottle choice, Sacc=saccharin

Female mice were then paired with drug naïve male mice while on the C2BC, until pregnancy was confirmed. A criteria of inclusion during the C2BC paradigm was for mice to drink above 10 mg/kg morphine during pregnancy, which has been shown to produce analgesia in rodents (Frances, Gout, Monsarrat, Cros, & Zajac, 1992; Fujita-Hamabe et al., 2012). As shown in Figure 2c, female mice continued to drink morphine in the C2BC throughout gestation and until their offspring reached PND 7, at which point pups were cross-fostered to a drug-naïve dam. During weeks 4-6 of the C2BC paradigm, dams displayed slightly lower morphine intake compared to week 2 and week 3 of the paradigm (RM mixed effects analysis; *F*(1.748, 17.19)= 4.807, *p* = 0.0256; Figure 2c). This phenomenon could be due to being paired with the male breeder for 5 hours during the day (week 4), and also to the increased bodyweight during pregnancy (week 5 and 6). Together, our data show that morphine-exposed dams exhibit signs of dependence upon removal of the drug and continue morphine drinking during pregnancy.

### 3.2 Neonate deficits before and after cross-fostering in offspring from morphine-exposed dams

To investigate the effects of maternal morphine exposure on offspring, pups were examined during early PNDs (Figure 1b). PND 3 offspring that were exposed to morphine through lactation displayed increased spontaneous activity (t-test; *t* = 2.527, *df* = 23, *p* = 0.0188; Figure 3a). Due to limited motor function during this early developmental period, the spontaneous signs monitored included audible cries, rolling over, and full 360° body rotation (Barr et al., 2011; Zhu & Barr, 2004).

**Figure 3:**
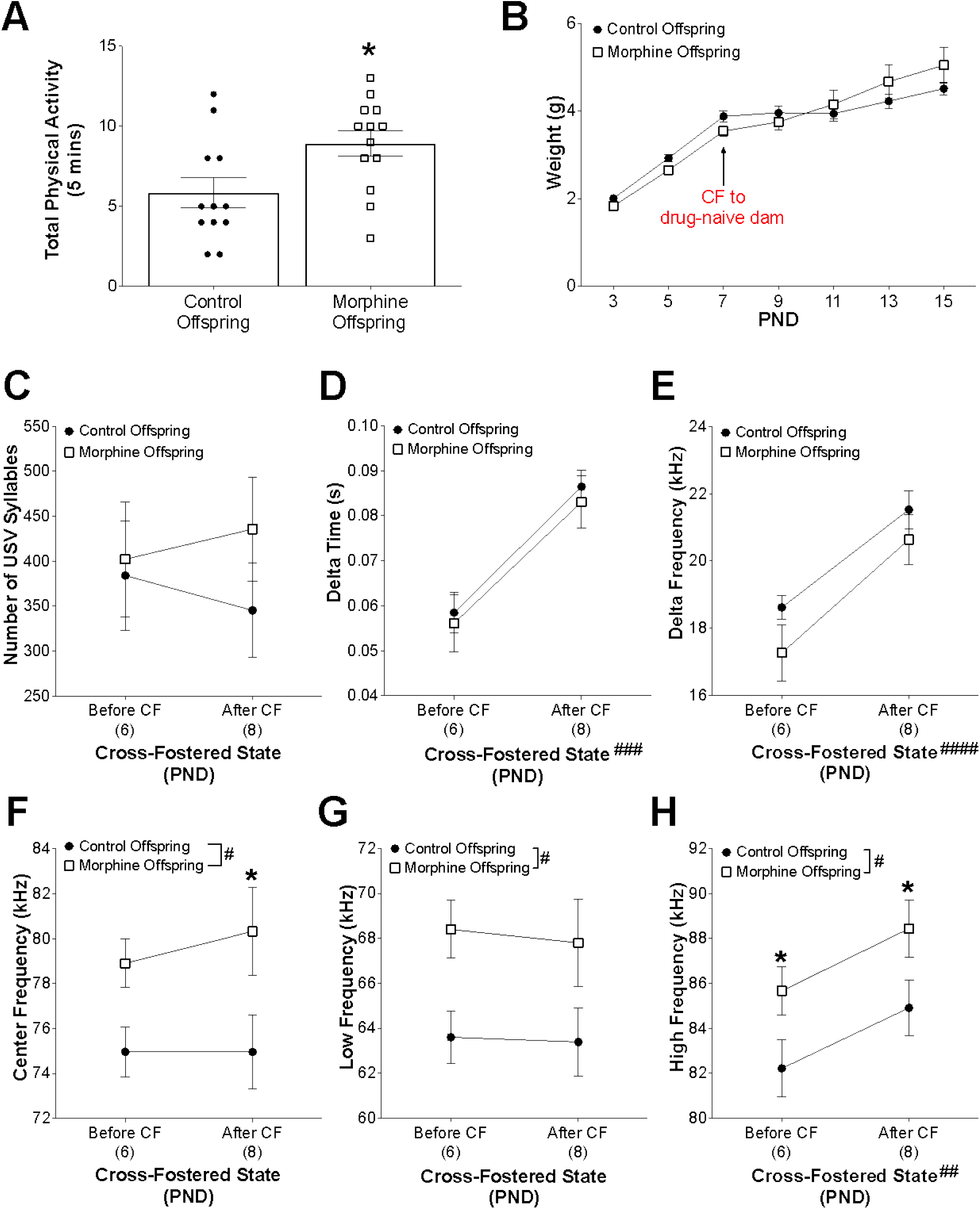
Behavioral outcomes in PND 2-15 pups before and after cross-fostering. **A**. PND 3 morphine offspring exhibited greater physical signs than control offspring (n=12,13). **B**. Offspring body weight before and after cross-fostering (PND 3-15); (10-18 litters/PND). **C**. Average number of USV syllables before and after CF (litter n= 9,7). **D**. Average delta time (s), or time duration, of each USV call (n= 9,7). **E**. Average delta frequency (Hz), or frequency range of each USV call (litter n= 9,7). **F**. Average center frequency (Hz), or middle frequency for each USV call (litter n= 9,7). **G**. Average low frequency (Hz) for each USV call (litter n= 9,7). **H**. Average high frequency (Hz) for each USV call (litter n= 9,7). PND= postnatal day, CF= Cross-Fostered. # indicates main effect: # p<0.05, ## p<0.01, ### p<0.001, #### p<0.0001; * indicates post-hoc significance: * p<0.05

To evaluate the long-term consequences associated with early-development morphine exposure, pups were cross-fostered to a drug-naïve dam on PND 7. This allowed for offspring to experience morphine withdrawal without introducing the dam’s drug-associated withdrawal behavior as a confound. Pups were weighed before and after cross-fostering, and an interaction of dam treatment x PND (RM mixed effects analysis; *F*(6, 160)=2.884, *p* = 0.0107) and a main effect of PND (RM mixed effects analysis; *F*(1.587, 42.31)=80.41, *p* = <0.0001) were observed (Figure 3b). Interestingly, morphine offspring had a trend for decreased body weight in early PNDs, compared to control offspring.

To evaluate distress that might be comparable to high-pitched crying seen in newborns with NOWS, ultrasonic vocalizations (USVs) were recorded in mice offspring before and after cross-fostering. We were specifically interested in evaluating USVs at PND 6 and PND 8, which corresponds to timepoints right before and after cross-fostering, respectively. This approach was chosen to evaluate changes while the offspring were lactating from morphine-treated dams (PND 6) and when they would potentially be undergoing acute drug withdrawal (PND 8) since they could no longer lactate from their respective dam. Offspring from morphine-exposed dams displayed no changes in the number of calls compared to control (Figure 3c). There was also no effect of dam treatment on delta time (i.e. length) of calls, but there was a main effect of PND (RM two-way ANOVA; *F*(1, 14)=18.22, *p* = 0.0008; Figure 3d). Similarly, there was no effect of dam treatment on the frequency range (delta frequency) of calls, but there was a main effect of PND (RM two-way ANOVA; *F*(1, 14)=39.70, *p* = <0.0001; Figure 3e).

However, there was a significant main effect of dam treatment on the frequency parameters of offspring’s USVs (Figure 3f-h). Offspring from morphine-exposed dams had calls of higher center frequency (RM two-way ANOVA; *F*(1, 14)=6.304, *p* = 0.0249; Figure 3f) compared to control offspring, and post-hoc analysis revealed that this change was statistically significant after cross-fostering (PND 8). Offspring from morphine-exposed dams also had calls of higher low-frequency points (RM two-way ANOVA; *F*(1, 14)=5.696, *p* = 0.0317; Figure 3g). In addition, offspring exposed to *in utero* morphine had higher high-frequency points in the calls (RM two-way ANOVA; *F*(1, 14)=4.713, *p* = 0.0476; Figure 3h), and there was a main effect of PND (RM two-way ANOVA; *F*(1, 14)=14.08, *p* = 0.0021; Figure 3h). Post-hoc analysis revealed that offspring from morphine-exposed dams had higher high-frequency points in their calls both before cross-fostering (PND 6) and after cross-fostering (PND 8), compared to control offspring. Overall, these results show that maternal morphine exposure alters neonatal spontaneous activity, body weight, and ultrasonic vocalization acoustic parameters.

### 3.3 Changes in anxiety-like/compulsive-like behavior in adolescent offspring from morphine-exposed dams

Offspring from morphine-exposed dams were evaluated for changes in behavior during adolescence to further understand the consequences of maternal opioid exposure during a critical period of development (Figure 4a). We used various behavioral tests to assess baseline changes in somatic and affective behavior, including measures to investigate locomotion, compulsive-like, anxiety-like, and depressive-like behavior. Behavioral measures were assessed for an effect of dam treatment and/or sex, but because no effect of sex was observed, males and females were combined. There was no difference in baseline physical signs between offspring from morphine-exposed dams and control offspring (Figure 4b). However, offspring from morphine-exposed dams buried more marbles than offspring from control dams in the marble burying test (MBT) (t-test; *t* =2.971, *df* = 69, *p*=0.0041; Figure 4c), displaying more anxiety-like/compulsive-like behavior.

**Figure 4:**
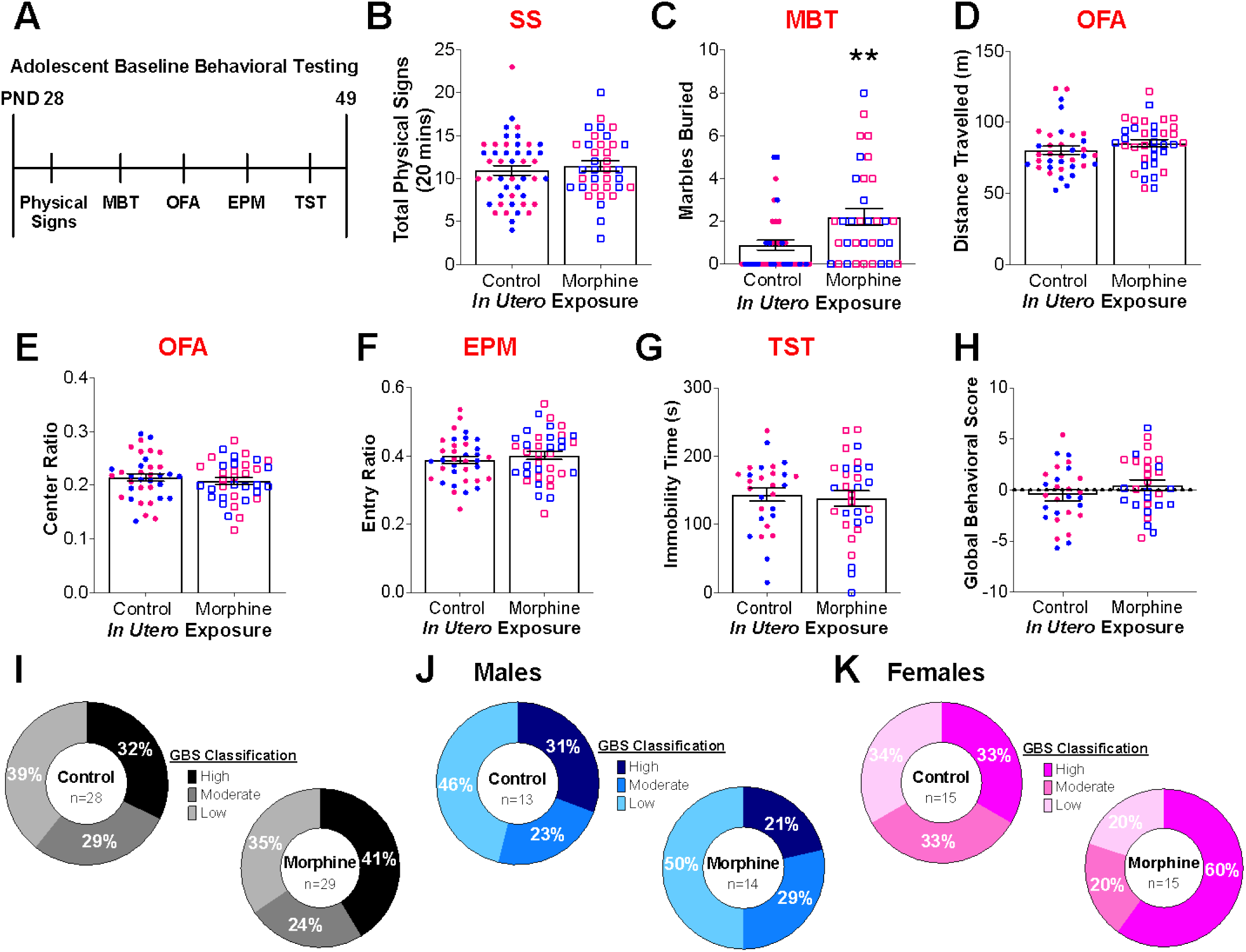
Sex-specific changes in baseline behavior in adolescent offspring from morphine-exposed dams. **A**. Experimental scheme showing the sequence of behavioral tests for the analysis of PND 28-49 adolescent offspring. **B**. Average number of total physical signs in control and morphine-exposed offspring (n=46, 36). **C**. Average number of marbles buried in the Marble Burying Test (MBT) (n=37, 34). **D**. Average distance travelled (m) in the Open Field Arena (OFA) (n=36, 36). **E**. Average center distance ratio in the OFA (n=36, 36). **F**. Average open arm entry ratio in the Elevated Plus Maze (EPM) (n=36, 37). **G**. Average immobility time (s) in the Tail Suspension Test (TST) (n=28, 30). **H**. Average global behavioral score (GBS) calculated as the summation of all the z-scores for each behavioral test for each animal (n=28, 29). Percent of offspring from control and morphine-exposed dams (**I)**, and male (**J)** and female **(K**) offspring that classified as high, moderate, and low scorers based on their global behavioral score for baseline behavior. **p<0.01 PND = postnatal day; SS = somatic signs; Blue symbols = males; Pink symbols = females

There was no effect of dam treatment in the Open Field Arena (OFA) for total distance travelled or center distance ratio (Figure 4d-e), nor did we detect significant differences in the open arm entry ratio in the Elevated Plus Maze (EPM; Figure 4f). Similarly, when offspring were assessed for depressive-like behavior in the Tail Suspension Test (TST), no significant difference in total immobility time was observed (Figure 4g).

Although there were no significant differences in various behaviors when individual tests were considered, we were interested in integrating multiple behavioral outcomes into a composite score. This would allow us to characterize offspring behavior holistically, which has been used in multiple areas of research (El-Kordi et al., 2013; Guyenet et al., 2010; Möller et al., 2018; O’Neal, Nooney, Thien, & Ferguson, 2020; Pereira de Souza Goldim et al., 2020; Shahi, Freedman, Dahl, Karandikar, & Mangalam, 2019). The use of a global severity score classification system allows us to examine the distribution of animals’ performance across multiple behavioral tests, where higher values represent higher behavioral symptom severity. As shown in Figure 4h, offspring from morphine-exposed dams have similar global behavioral scores (GBS) compared to offspring from control dams. To determine the distribution of adolescent offspring GBS, experimental scores were characterized into ‘high’ (GBS>1), ‘moderate’ (1<GBS<1), and ‘low’ (GBS<1) phenotypes. Because there was a trend (p=0.0898) for a main effect of dam treatment when the three GBS classifications were evaluated for the offspring (data not shown), we evaluated the percentage of offspring in each GBS classification. Among control offspring (n=28), 39% were classified as having a ‘low’ GBS phenotype, 29% were ‘moderate’, and 32% were ‘high’ (Figure 4i). However, offspring from morphine-exposed dams (n=29) had a higher percentage of ‘high scores’ (41%), and 35% classified as having a ‘low’ severity phenotype, while 24% were ‘moderate’ (Figure 4i). In addition, when sex was investigated, 46%, 23%, and 31% of male control offspring fell under the ‘low’, ‘moderate’, and ‘high’ classification, respectively (Figure 4j). Male offspring from morphine-exposed dams were characterized at similar percentages in each GBS classification (low=50%, moderate=29%, and high=21%) (Figure 4h). Conversely, female offspring from morphine-exposed dams had a trend for a higher percentage being classified in the ‘high’ category (60%), compared to control female offspring (33%) (Figure 4k). Twenty percent of female offspring from morphine-exposed dams were categorized as ‘moderate’ and ‘low’ scorers based on their GBS, while 33%-34% of control female offspring were categorized as ‘moderate’ and ‘low’ (Figure 4k). Although the GBS classification is not significantly different between offspring, the higher percent of ‘high’ GBS phenotype in the offspring from morphine-exposed dams is intriguing in that it suggests that early life morphine exposure might lead to an increase in the number of offspring that have a more severe phenotype when considering a broad array of behaviors, as opposed to very significant deficits in any one behavioral measure.

### 3.4 Changes in baseline behavior in adult offspring from morphine-exposed dams

We were interested in the possibility that the behavioral phenotypes we observed could persist beyond adolescence and into adulthood. Therefore, in a separate cohort of control and morphine-exposed offspring, we evaluated baseline adult behavior to determine the long-term effects of maternal morphine exposure using the same battery of behavioral tests used for adolescent mice (Figure 5a). Behavioral measures were assessed for an effect of dam treatment and/or sex, but when no effect of sex was detected, males and females were combined. Offspring from morphine dams did not significantly differ from control offspring in baseline total physical signs (Figure 5b), compulsive-like behavior in the MBT (Figure 5c), locomotion or anxiety-like behavior in the OFA (Figure 5d-e), and depressive-like behavior in the TST (Figure 5h). However, when adult offspring were evaluated for anxiety-like behavior in the EPM, offspring from morphine-exposed dams displayed no difference in entry ratio (Figure 5g), but did display decreased time spent in the open arms, compared to control offspring (t-test; *t* = 2.935, *df* = 35, *p* = 0.0059; Figure 5f). This suggests that offspring from morphine-exposed dams have increased anxiety-like behavior in adulthood.

**Figure 5:**
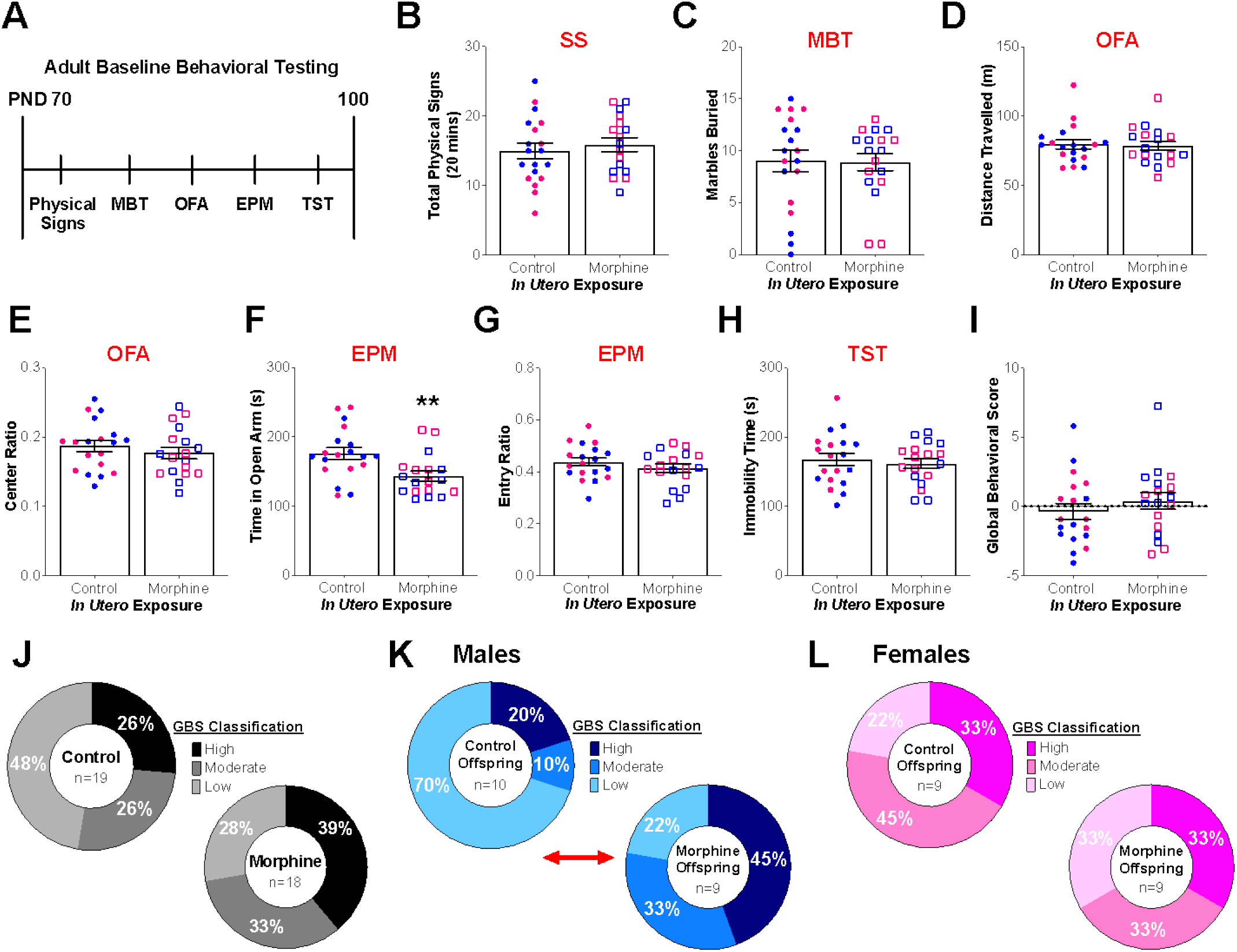
Sex-specific changes in baseline behavior of adult offspring from morphine-taking dams. **A**. Experimental scheme for the analysis of adult offspring in a battery of behavioral tests. **B**. Average baseline total physical signs observed in offspring (n=19, 17). **C**. Average marbles buried in the Marble Burying Test (MBT) (n=19, 18). **D**. Average distance travelled (m) in the Open Field Arena (OFA) (n=19, 18). **E**. Average center distance ratio in the OFA (n=19, 18). **F**. Average time in the open arms of the Elevated Plus Maze (EPM) (n=19, 18). **G**. Average open arm entry ratio in the EPM (n=19, 18). **H**. Average immobility time (s) in the Tail Suspension Test (TST) (n=19, 18). **I**. Average global behavioral scores (GBS) calculated as the summation of all the z-scores for each behavioral test for each animal (n=19, 18). Percent of control and morphine-exposed offspring (**J**), male (**K)**, and female (**L**) offspring that classified as high, moderate, and low scorers based on their GBS for baseline behavior. **p<0.01; SS = somatic signs; Blue symbols = males; Pink symbols = females

We also used the GBS to integrate the multiple behavioral outcomes into a composite score which allowed us to characterize adult offspring behavior holistically, as described above. Although there was no difference in overall global behavioral score between offspring from morphine-exposed dams and control dams (Figure 5i), the percentage of offspring that fell into each GBS classification was of interest. For control offspring (n=19), 48% were classified as having a ‘low’ severity phenotype, 26% were ‘moderate’, and 26% were ‘high’ (Figure 5j). However, among offspring from morphine-exposed dams (n=18), only 28% were classified as ‘low’, 33% were ‘moderate’, and 39% were ‘high’ (Figure 5j), suggesting that a higher percentage of offspring from morphine-exposed dams might be more behaviorally vulnerable. Although the sample size was small, Figure 5k shows that the percentage of male offspring from morphine-exposed dams in each GBS classification was different than that of male control offspring (chi-square test; DF=2; *p* =0.0052). Among male control offspring (n=10), 70% were classified as having a ‘low’ GBS severity phenotype, 10% were ‘moderate’, and 20% were ‘high’ (Figure 5k). However, among male offspring from morphine-exposed dams (n=9), 22% were ‘low’, 33% were ‘moderate’, and 45% were ‘high’ (Figure 5k). Conversely, female offspring from morphine-exposed dams had a similar percent of offspring that fell into the three GBS classifications when compared to female control offspring (Figure 5l). For example, 33% of mice were classified as having a ‘high’ GBS severity phenotype in both groups.

Together, our results suggest that maternal morphine exposure has long-term consequences throughout the offspring’s life span, as reflected by changes that persist in adulthood. Offspring from morphine-exposed dams display increased baseline levels of anxiety-like behavior during adulthood. In addition, a much higher percent of male offspring from morphine-exposed dams fall into the high and moderate GBS severity classification. This suggests that not only are specific phenotypes altered by *in utero* opioid exposure, but that, overall, male offspring are at risk of developing more severe behavioral phenotypes during adulthood, a phenomenon that could be revealed or exacerbated by stress or drug use.

### 3.5 Male offspring from morphine-exposed dams display decreased two-hour ethanol intake

Given the well-documented interactions between alcohol and the opioid system (Arias & Kranzler, 2008; Berrettini, 2013; Gianoulakis, 2001; Gianoulakis et al., 1989; Job et al., 2007; Oslin, Berrettini, & O’Brien, 2006), we next wanted to assess alcohol use risk in offspring maternally exposed to morphine. The offspring tested in the battery of behavioral tests during adolescence were allowed to mature into adulthood and were then evaluated in an ethanol intermittent two-bottle choice (I2BC) paradigm (Figure 6a), which has been used to assess voluntary ethanol intake (Carnicella et al., 2014; Hwa et al., 2011; Quijano Cardé & De Biasi, 2022; Quijano Cardé et al., 2021). A three-way ANOVA revealed a main effect of sex where female mice (regardless of treatment) drank significantly more ethanol than male mice, so data and analyses are presented separately for each sex. Alcohol-related behaviors were evaluated at three different phases of the I2BC – Acquisition, Maintenance, and Fading - for both male and female offspring.

**Figure 6:**
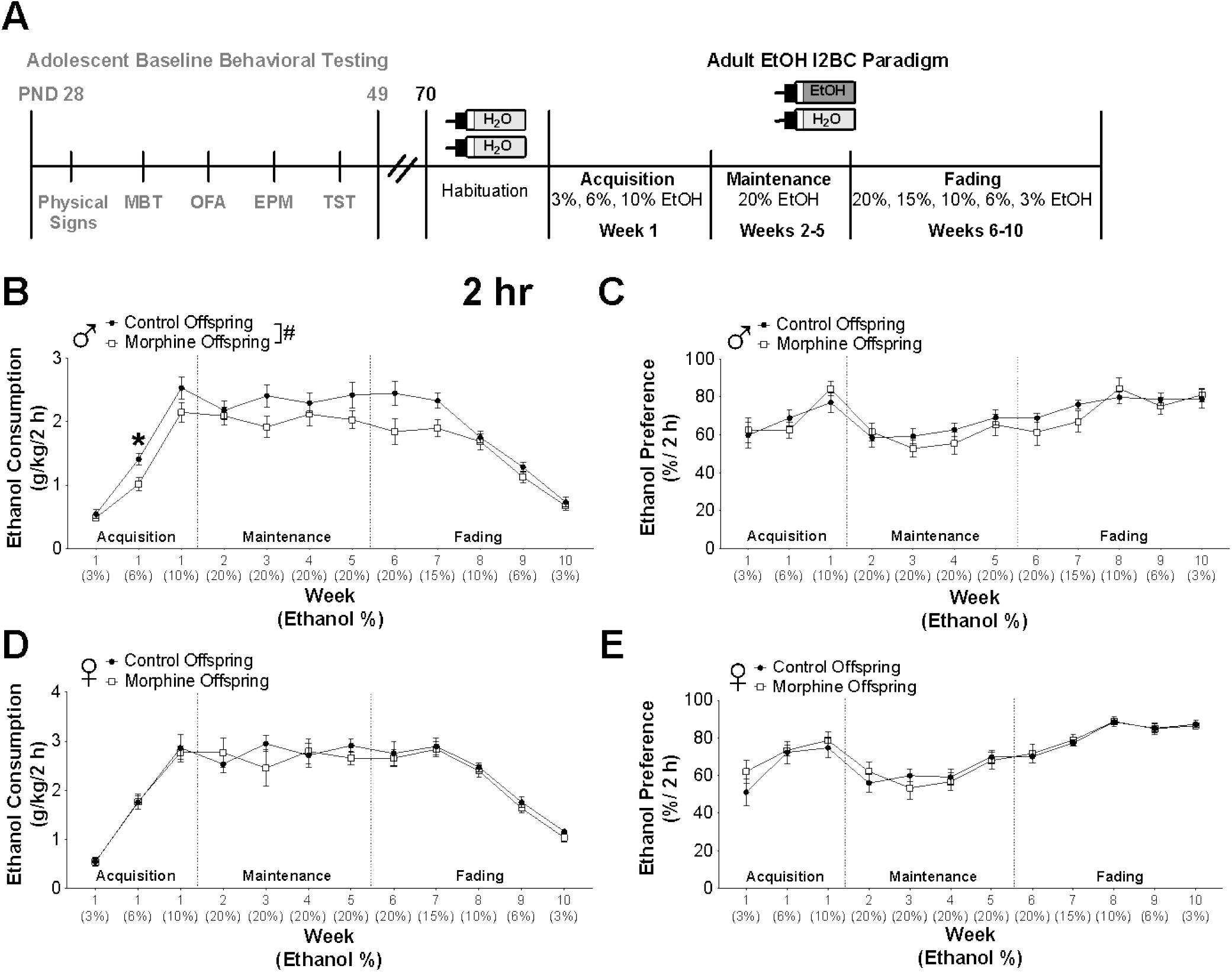
Ethanol 2-hour intake and preference for male and female offspring in the I2BC paradigm. **A** Schematic of the adult ethanol I2BC experimental timeline after mice underwent baseline adolescent behavioral testing. (**B & C**) Average 2-hr ethanol intake (g/kg) (B) and preference (%) (C) for male offspring during weeks 1-10 of ethanol drinking (n=14). **(D & E)** Average 2-hr ethanol intake (g/kg) (D) and preference (%) (E) for female offspring during weeks 1-10 of ethanol drinking (n=12). # p<0.05 main effect of dam treatment for ‘Acquisition’ and ‘Fading’ phase; * indicates post-hoc significance: * p<0.05 PND = postnatal day; MBT = marble burying test; OFA = open field arena; EPM = elevated plus maze; TST = tail suspension test; I2BC = intermittent two-bottle choice

#### Acquisition phase of I2BC

Ethanol drinking patterns were first evaluated for the ‘Acquisition’ phase of the I2BC, where mice were given increasing concentrations of ethanol during the first week of exposure. Figure 6b shows a main effect of concentration for 2-hour intake during the ‘Acquisition’ phase for male offspring (RM mixed effects analysis; *F*(1.417, 34.02) = 133.9, *p* <0.0001) and a main effect of dam treatment (RM mixed effects analysis; *F*(1, 26) = 6.176, *p*=0.0197). Specifically, post-hoc analysis revealed that male offspring from morphine-exposed dams drink less ethanol (g/kg) at the 6% concentration during the first two hours of the session, compared to male control offspring. When ethanol intake (g/kg) was evaluated during the 24-hour sessions of the ‘Acquisition’ phase (Supplemental figure 1a), although not significant, a trend (p=0.0791) for an effect of dam treatment was present for male offspring. There was a main effect of concentration for the 24-hour ethanol intake (RM two-way ANOVA; *F*(1.648, 42.86) = 138.9, *p* < 0.0001; Supplemental figure 1a). With regards to male offspring’s 2-hour ethanol preference (%) during the ‘Acquisition’ phase (Figure 6c), there was a main effect of concentration (RM mixed effects analysis; *F*(1.801, 43.22) = 8.540, *p* <0.0001), but no main effect of dam treatment. In addition, there was no main effect or interaction of concentration and/or dam treatment during male offspring’s 24-hour ethanol preference during the ‘Acquisition’ phase (Supplemental figure 1b).

In female offspring, the 2-hour session for the ‘Acquisition’ phase of the I2BC revealed a significant main effect of concentration for ethanol intake (g/kg) (RM mixed effects analysis; *F*(1.770, 35.39) = 134.3, *p* <0.0001; Figure 6d) and preference (%) (RM mixed effects analysis; *F*(1.621, 32.42) = 7.813, *p* = 0.0030; Figure 6e), but no effect of dam treatment. Similarly, for female offspring’s 24-hour ethanol intake (g/kg) (Supplemental figure 1c) there was a main effect of concentration (RM two-way ANOVA; *F*(1.818, 40.00) = 175.6, *p* < 0.0001), but no effect of dam treatment. Although not significant, there was a trend (p=0.0699) for a main effect of concentration, but no effect of dam treatment on female offspring’s 24-hour ethanol preference for the ‘Acquisition’ phase (Supplemental figure 1d).

Together, this reveals that male -but not female-offspring from morphine-exposed dams drink lower amounts of ethanol during the ‘Acquisition’ phase of the I2BC, but have no changes in ethanol preference during this part of the experimental paradigm.

#### Maintenance phase of I2BC

Ethanol drinking patterns were next evaluated during the ‘Maintenance’ phase of the I2BC, where mice were exposed every other day for four weeks (weeks 2-5) to two bottles, one containing 20% ethanol and the other containing water. In male offspring during the ‘Maintenance’ phase, there was no effect of week, and although not significant, a trend (p=0.0793) was present for dam treatment for two-hour ethanol intake (g/kg) (Figure 6b). In addition, there was no effect of week or dam treatment for males’ 24-hour ethanol intake (g/kg) during the ‘Maintenance’ phase (Supplemental figure 1a). With regards to male offspring’s ethanol preference (%) during the ‘Maintenance’ phase, there was a main effect of week for the two-hour session (RM two-way ANOVA; *F*(2.163, 56.23) = 3.369, *p*= 0.0380; Figure 6c) and 24-hour session (RM two-way ANOVA; *F*(1.900, 49.40) = 6.162, *p*= 0.0047; Supplemental figure 1b), but no main effect of dam treatment.

Female offspring did not show a significant difference of week or dam treatment for 2-hour (Figure 6d) and 24-hour (Supplemental figure 1c) ethanol intake (g/kg) during the ‘Maintenance’ phase of the I2BC. However, there was a main effect of week for both 2-hour (RM mixed effects analysis; *F*(2.746, 59.50) = 4.141, *p* = 0.0118; Figure 6e) and 24-hour ethanol preference (RM two-way ANOVA; *F*(2.848, 62.66) = 6.698, *p* = 0.0007; Supplemental figure 1d), but no effect of dam treatment.

Together, our data show that ethanol intake and preference during the 20% ethanol ‘Maintenance’ phase of the I2BC in both male and female offspring from morphine-exposed dams are similar to control, implying that there is no effect of dam treatment.

#### Fading phase of I2BC

Lastly, ethanol drinking patterns were evaluated for the ‘Fading’ phase of the I2BC, where mice were given decreasing concentrations of ethanol (20%, 15%, 10%, 6%, 3%) for the remaining five weeks of the paradigm. As shown in Figure 6b, analysis of the 2-hour ethanol intake during the ‘Fading’ phase for male offspring revealed a main effect of concentration (RM two-way ANOVA; *F*(1.960, 50.95) = 79.10, *p* <0.0001), a main effect of dam treatment (RM two-way ANOVA; *F*(1, 26) = 4.267, *p*=0.0490), and a concentration x dam treatment interaction (RM two-way ANOVA; *F*(4, 104) = 3.079, *p*=0.0193). Specifically, our data suggest that male offspring from morphine-exposed dams consume less ethanol at various concentrations during the 2-hour ‘Fading phase’ of the paradigm. When the 24-hour ethanol intake (g/kg) during the ‘Fading’ phase was evaluated, there was a main effect of concentration (RM two-way ANOVA; *F*(2.562, 66.62) = 142.4, *p* < 0.0001; Supplemental figure 1a), but not dam treatment. With regards to male offspring’s 2-hour ethanol preference (%) during the ‘Fading’ phase, there was a main effect of concentration (RM two-way ANOVA; *F*(2.335, 60.71) = 11.19, *p* <0.0001; Figure 6c), but no main effect of dam treatment and a trend for an interaction (p=0.0927). Similarly, during male offspring’s 24-hour ethanol preference in the ‘Fading’ phase, there was a main effect of concentration (RM two-way ANOVA; *F*(2.155, 56.04) = 84.08, *p* <0.0001; Supplemental figure 1b), but no main effect of dam treatment.

No differences were detected when comparing control and morphine-exposed female offspring. We found a significant main effect of concentration for ethanol intake (g/kg) (RM two-way ANOVA; *F*(2.219, 48.81) = 72.56, *p* <0.0001; Figure 6d) and preference (%) (RM two-way ANOVA; *F*(2.664, 58.62) = 24.01, *p* <0.0001; Figure 6e) at the 2-hour timepoint during the ‘Fading’ phase of the I2BC, but no effect of dam treatment. Similarly, at 24-hour there was a significant main effect of concentration for ethanol intake (g/kg) (RM two-way ANOVA; *F*(2.049, 45.07) = 108.6, *p* <0.0001; Supplemental figure 1c) and preference (%) (RM two-way ANOVA; *F*(3.099, 68.19) = 144.0, *p* <0.0001; Supplemental figure 1d), but no effect of dam treatment.

Overall, our results indicate that male -but not female-offspring from morphine-exposed dams drink lower amounts of ethanol during the initial 2-hour phase of the ‘Fading’ experiment although there are no changes in ethanol preference.

It should be noted that there was a main effect of dam treatment (RM two-way ANOVA; *F*(1,26) = 7.678, *p* = 0.0102; Supplemental Figure 2b) for 24-hour total fluid intake for male offspring, and a main effect of week (RM two-way ANOVA; *F*(2.576,66.98) = 18.00, *p* = <0.0001), where male offspring from morphine-exposed dams had lower total fluid intake compared to male control offspring. This main effect of dam treatment for total fluid intake was not observed at the two-hour timepoint (Supplemental Figure 2a, 2c).

## 4 Discussion

Our maternal morphine C2BC paradigm demonstrated that morphine dams display signs of dependence and voluntarily drink morphine throughout gestation. Maternal morphine exposure with this paradigm increases neonate spontaneous activity, decreases body weight before cross-fostering, and alters various USV frequency parameters. The set of experiments presented in this study also demonstrates subtle sex-specific alterations in adolescent and adult offspring exposed to pre- and perinatal morphine. Overall, the data presented supports the hypothesis that maternal opioid exposure alters offspring behavior throughout development.

Our study is one of few to investigate offspring outcomes using a maternal oral self-administration model that starts before gestation, continues throughout gestation, and extends one week postnatally. Most preclinical studies investigating the effects of *in utero* morphine exposure on offspring behavior have used daily injections or forced oral solution, making it difficult to discern whether the effect seen in offspring is due to an interaction of maternal stress with opioid exposure, or solely due to opioid exposure (Chiou et al., 2003; Glick et al., 1977; Klausz et al., 2011; Nasiraei-Moghadam et al., 2013; Siddiqui et al., 1997; Sobor et al., 2010; Timár et al., 2010; P. L. Wu et al., 2018; Yang et al., 2003). In addition, the duration of maternal opioid administration varies across studies. For example, studies differ among each other for using a partial gestation, full gestation, or gestation and lactation maternal opioid paradigm (Chiou et al., 2003; De Vries et al., 1991; Eriksson & Rönnbäck, 1989; Gagin et al., 1997; Glick et al., 1977; Jóhannesson & Becker, 1972; Klausz et al., 2011; Laborie et al., 2005; Ramsey et al., 1993; Rimanóczy, Ŝlamberová, Riley, & Vathy, 2003; Schindler et al., 2004; Shen et al., 2016; Tan et al., 2015; Timár et al., 2010; P. L. Wu et al., 2018; Yang et al., 2003). Each of these exposure paradigms models specific critical developmental periods for the fetus and can confer paradigm-specific behavioral alterations.

An extended maternal morphine treatment that continues throughout the lactation period might be necessary to recapitulate the clinical outcomes of prenatal morphine and NOWS, considering that rodent gestation/early offspring postnatal period has been compared to late human gestation and newborn birth, when considering various morphological and functional milestones relating to eye, cardiac, immune, and brain development (Clancy, Darlington, & Finlay, 2001; Craig et al., 2003; Holsapple, West, & Landreth, 2003; Krishnan et al., 2014; Kroon, van Hugte, van Linge, Mansvelder, & Meredith, 2019; Lazic, 2012; Rice & Barone, 2000; Richard & Flamant, 2018; Van Cruchten et al., 2017). Due to the short gestation period compared to humans, many developmental processes (e.g. myelination and immune function) continue postnatally in rodents (Craig et al., 2003; Holsapple et al., 2003). Caution is therefore warranted when making direct developmental comparisons between species since this is dependent on the processes being investigated and the developmental window studied. Overall, early rodent postnatal days might be an important consideration when developing maternal drug exposure paradigms. For this reason, we cross-fostered the offspring at PND 7 to a drug-naïve dam rather than removing the morphine bottle from the dam, thereby preventing maternal withdrawal behavior from becoming a confound in the study. We also wanted the rodent pups to undergo morphine withdrawal and potentially experience characteristics of NOWS that can be investigated before weaning and might produce long-term behavioral consequences. Most clinical and preclinical studies have shown that offspring from opioid-dependent mothers display either reduced body weight or no change in body weight (Corr, Schaefer, & Paul, 2018; Dutriez-Casteloot et al., 1999; Gagin et al., 1997; Jones et al., 2010; Kaltenbach et al., 2018; Klausz et al., 2011; Laborie et al., 2005; Ramsey et al., 1993; Shen et al., 2016; Siddiqui et al., 1997; Siu & Robinson, 2014; Timár et al., 2010). However, some preclinical studies like Chiang *et al*. (2010) and Timar *et al*. (2010) have reported increased body weight in PND 7, PND 14, and PND 21 offspring after maternal morphine exposure. Although we anticipated decreased body weight in offspring from morphine-exposed dams even after cross-fostering, our data suggest that cross-fostering increases overall pup mortality (data not shown) and might stunt body weight gain in control offspring. This phenomenon has also been reported in other models of cross-fostering (Santangeli et al., 2016). In addition, cross-fostering has been shown to affect both maternal and offspring behavior (Dulor Finkler, Espinoza Pardo, & Bolten Lucion, 2020; Gauthier, Deangeli, & Bucci, 2015; R. Šlamberová, Hrubá, Bernášková, Matějovská, & Rokyta, 2010; I. Vathy, Šlamberová, & Liu, 2007), likely due to the stress associated with the new environment and alterations in the mother-infant relationship. This posits the question of whether the drug-naïve dam euthanized the most “vulnerable” offspring, and whether the relatively subtle behavioral effects we observed between control and morphine-exposed offspring might be due to the fact that we tested the “resilient” offspring that survived after cross-fostering.

Preclinical studies aim to model maternal opioid exposure that results in offspring outcomes comparable to those of human newborns experiencing NOWS. For example, studies have examined developmental milestones in rodents, as well as pup mortality and bodyweight and compared them to newborn outcomes in clinical studies (Chiang, Hung, Lee, Yan, & Ho, 2010; Dutriez-Casteloot et al., 1999; Eriksson & Rönnbäck, 1989; Gagin et al., 1997; Jóhannesson & Becker, 1972; Laborie et al., 2005; Ramsey et al., 1993; Siddiqui et al., 1997; Sobor et al., 2010; Timár et al., 2010). To our knowledge, this is the first study to investigate how <u>maternal morphine</u> exposure alters ultrasonic vocalizations in pups, as a correlate to high-pitched crying in human newborns and as a characteristic of NOWS. Similar to our results, one study found that offspring from oxycodone-exposed dams have higher frequency USVs than control offspring (Zanni et al., 2021). Another study also found that neonate offspring injected with morphine from PND 1 – 14 had increased USV frequency parameters (Borrelli et al., 2021). Various studies and reviews have focused on understanding vocalizations and changes in their acoustic parameters, including how upward shifts in frequency modulation is usually associated with an increase in infant distress (Brudzynski, 2015; Castellucci, Calbick, & McCormick, 2019; Esposito, Nakazawa, Venuti, & Bornstein, 2013; Hahn & Lavooy, 2005; Kromkhun et al., 2013; Lingle, Wyman, Kotrba, Teichroeb, & Romanow, 2012; Parga et al., 2020; Wasz-Höckert, Michelsson, & Lind, 1985). Although our work did not further probe the specific brain-regions and mechanisms that lead to alterations in USV frequency parameters in offspring from morphine-exposed dams, other studies have shown that the periaqueductal grey (PAG), a brain region with dense expression of opioid receptors, and the opioid receptor system in general, are important for USV syllable production (D’Amato, 2021; Goodwin & Barr, 2005; Tschida et al., 2019). For example, PAG lesions decrease USVs in pups (Wiedenmayer, Goodwin, & Barr, 2000) and mu-opioid receptor knockout (*Orpm*^−/−^) pups emit less USV calls compared to their littermates in response to maternal isolation (Moles, Kieffer, & D’Amato, 2004). Offspring exposed to opioids *in utero* display changes in the opioid receptor system (Chiou et al., 2003; Ilona Vathy, Šlamberová, Rimanóczy, Riley, & Bar, 2003), which further supports our finding that offspring from morphine-exposed dams have profound changes in USV-related parameters. The functional role of brain-region specific changes in opioid receptor and endogenous opioid levels in areas such as the PAG merits further investigation.

In addition to changes in neonatal outcomes, offspring from morphine-exposed dams also display changes in baseline behavior during adolescence and adulthood. Although we found no significant differences in locomotion or depressive-like behavior, adolescent offspring from morphine-exposed dams displayed increased anxiety-like/compulsive-like behavior in the MBT. During adulthood, offspring from morphine dams displayed increased anxiety-like behavior in the EPM. Contrary to what we found, a few preclinical studies investigating the effects of prenatal morphine exposure found either decreased anxiety-like behavior or no changes in anxiety-related behaviors, which highlights how duration and dose of maternal morphine exposure can have seemingly opposite effects in offspring (Klausz et al., 2011; Tan et al., 2015). Interestingly, similar to our results of increased anxiety-like/compulsive-like behavior in offspring from morphine-exposed dams, male offspring from morphine-exposed parents displayed decreased percent open arm time in the EPM (Sabzevari et al., 2019; Vousooghi et al., 2018), increased grooming, and increased marbles buried (Rohbani et al., 2019), suggesting increased anxiety-like/compulsive-like behavior. Together with our results, these data suggest increased behavioral vulnerability in male offspring from morphine-exposed dams, while there are no apparent changes in female offspring behavior. In clinical studies, few have investigated how prenatal exposure to opioids affects males and females differently.

The variability observed in preclinical findings is likely attributable, among other factors, to the inherent vulnerability and/or resiliency of sub-populations of newborns, and to the many differences in maternal drug exposure paradigms. These differences include dose of drug, duration and timing of exposure, and route of drug administration. This not only leads to pharmacokinetic and pharmacodynamic differences but also differentially impacts the stress experienced by the dam during treatment, a phenomenon that may lead to changes in offspring behavior. Similar discrepancies are also a consideration for retrospective clinical studies and can make cross-study comparisons difficult.

Prenatal or early life stress can increase susceptibility to various behavioral manifestations in male rodents later in adulthood (Columba-Cabezas, Iaffaldano, Chiarotti, Alleva, & Cirulli, 2009; Lebow et al., 2019; Sarkar, 2015). In this context, prenatal opioid exposure and the experience of NOWS might also be viewed as a stressor capable of modifying behavior later in life. One potential explanation for observing changes in adulthood - and not adolescence - in male offspring could be that a more severe behavioral phenotype is unmasked among male offspring from morphine-exposed dams once all hormonal, chemical, and circuitry-related changes have matured in adulthood (Sinclair, Purves-Tyson, Allen, & Weickert, 2014). Interestingly, a higher percentage (45% vs. 20% in control offspring) of male offspring from morphine-exposed dams were classified as having higher and more severe global behavioral scores during adulthood. Merhar *et al*. (2019) reported that 40% of opioid-exposed newborns exhibit significant brain alterations, suggesting that the disruption of key processes during the development of the nervous system might increase vulnerability to behavioral deficits later in life. Similar to the percentage value in the clinical data, our study finds that 39% of adolescent offspring and 41% of adult offspring from morphine-exposed dams fall under the ‘High’ GBS classification, suggesting a more severe behavioral phenotype. Analyzing offspring behavior using a composite score might therefore help to identify vulnerable sub-populations of individuals that need additional non-pharmacological and/or pharmacological interventions At a minimum, such stratification might improve testing drug efficacy, as proposed by the Food and Drug Administration (FDA, 2019).

Offspring exposed prenatally to morphine have altered sensitivity to drugs, including morphine, cocaine, and methamphetamine, and display changes in drug-reward related behavior (Akbarabadi et al., 2018; Chiang et al., 2014; Gagin et al., 1997; Glick et al., 1977; He, Bao, Li, & Sui, 2010; Jiang, He, Wang, & Sui, 2011; Ramsey et al., 1993; Sadat-Shirazi et al., 2019; Shen et al., 2016; Timár et al., 2010; Wang, Yao, Li, Nie, & He, 2017; L. Y. Wu et al., 2009). Nygaard *et al*. (2020) showed that although there were no significant differences in alcohol consumption in a one-year span in adults whose mothers misused heroin, a significantly higher proportion of those individuals reported misusing alcohol during their lives. To date, no preclinical study seems to have established a relationship between *in utero* morphine exposure and offspring alcohol use. Although we hypothesized that offspring exposed to pre- and perinatal morphine would have higher ethanol intake and preference, we surprisingly found that male offspring from morphine-exposed dams had lower 2-hour ethanol intake, compared to male control offspring, despite no changes in alcohol preference. ‘Front-loading’ behavior, wherein the largest amount of ethanol consumed is observed toward the onset of EtOH access, is thought to reflect increased motivation to experience the rewarding effects of ethanol (Darevsky et al., 2019; Linsenbardt & Boehm, 2014; Rhodes et al., 2007; Salling et al., 2018; Wilcox et al., 2014), and, therefore, it is tempting to speculate that early exposure to morphine changes the subjective reward to ethanol. Among other mechanisms, alcohol leads to hypothalamic activation and increased levels of glucocorticoids which modify reward-related behaviors by stimulating mesencephalic dopaminergic transmission and increasing norepinephrine (NE) levels in the prefrontal cortex (PFC) (Piazza & Le Moal, 1997). Reduced drinking in morphine-exposed male offspring at the 2-hour timepoint might be due to hypoactivity of the stress response and/or hypothalamic-pituitary-adrenal (HPA) axis, which has been shown to be dysregulated in rodent offspring exposed to *in utero* morphine (Klausz et al., 2011; Laborie et al., 2005; Rimanóczy et al., 2003; Romana Šlamberová, Rimanóczy, Riley, & Vathya, 2004). Further studies are needed to understand the influence of maternal morphine exposure on HPA axis function, and consequently the effects on ethanol use. It is still unclear how alterations in fetal development by gestational opioids compound with other factors, such as stress or subsequent drug exposure, manifest in adulthood.

## 5 Conclusion

The data presented supports the hypothesis that prenatal-perinatal morphine exposure alters offspring behavior. Although not modeled in our study, important factors that influence and can exacerbate human offspring outcomes include: mother’s poly-drug use, socioeconomic stress experienced by pregnant mothers, and stressors experienced by the offspring. Questions left to be answered include whether or not stress during adolescence and/or adulthood can “push” offspring exposed prenatally to opioids from the ‘moderate’ behavioral severity phenotype to the ‘high’ category, and whether stress can alter adolescent and adult drug reward sensitivity, including commonly used drugs like ethanol and tobacco products.

## Data Availability Statement

The data that support the findings of this study are available from the corresponding author upon reasonable request.

## Funding Statement and Acknowledgements

This work was supported in part by NIH DA044205, DA049545, and U01 AA025931 to MDB, and NSF-GRFP grant DGE-1321851 to VCF. We thank Dr. Gordon Barr, Dr. Amelia Eisch, and Dr. Giulia Zanni for lending us the Dodotronic microphone and for their training in recording USVs and using the Raven software. We thank Melanie Schaffler for piloting the USV protocol, and Kimberly Halberstadter for partial collection of neonate and adult drinking data. We also thank Dr. Theresa Patten and Dr. Natalia A. Quijano Cardé for insightful feedback on this manuscript.

## Ethics Approval Statement

This work involved rodents in its research. The authors confirm that all animal subject research was conducted with the approval of the Institutional Animal Care and Use Committee (IACUC) at the University of Pennsylvania.

