## Supplemental Figures for "In utero exposure to morphine leads to sex-specific behavioral alterations that persist into adulthood in cross-fostered mice"

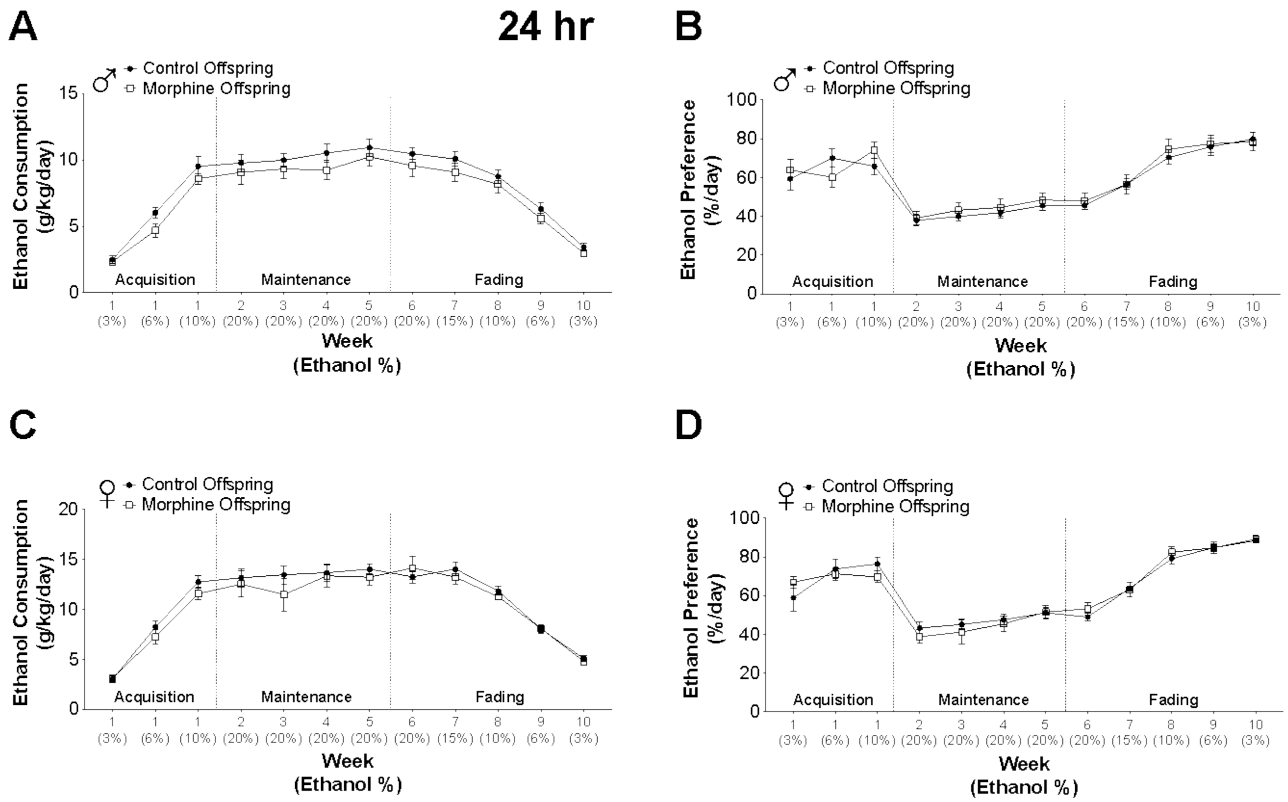


**Supplemental Figure 1**

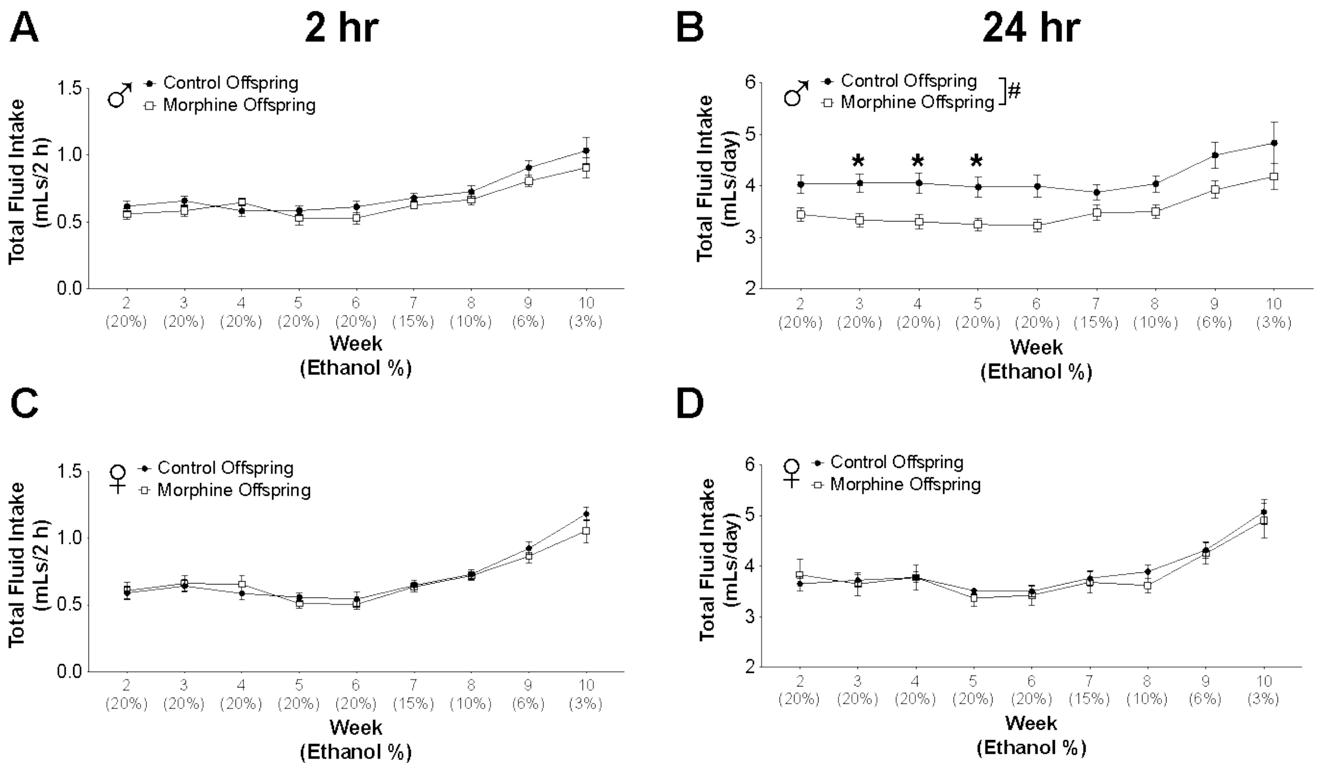


**Supplemental Figure 2**

**Figure legends:**

**Supplemental Figure 1**:

**Ethanol intake and preference (24-hour) for male and female offspring in the I2BC paradigm.**

**(A & B)** Average 24-h ethanol intake (g/kg) (A) and average 24-h ethanol preference (%) (B) for male offspring during weeks 1-10 of drinking (n=14). (**C & D)** Average 24-h ethanol intake (g/kg) (C) and ethanol preference (%) (D) for female offspring during weeks 1-10 of drinking (n=12).

I2BC = intermittent two-bottle choice

**Supplemental Figure 2**:

**Total fluid intake (2- and 24- hour) for offspring in the I2BC paradigm. A.** Average 2-hour total fluid intake (mL) for male offspring from weeks 2-10 (n=14). **B**. Average 24-hour total intake (ml) for male offspring from weeks 2-10 (n=14). **C**. Average 2-hour total fluid intake for female offspring from weeks 2-10 (n=12). **D**. Average 24-hour total fluid intake for female offspring from weeks 2-10 (n=12).

### p<0.05 main effect of dam treatment; * indicates post-hoc significance: * p<0.05
